# Partial-resistance against aphids in wild barley reduces the oviposition success of the generalist parasitoid, *Aphidius colemani* Viereck

**DOI:** 10.1101/653113

**Authors:** Daniel. J. Leybourne, Tracy. A. Valentine, Jorunn. I. B. Bos, Alison. J. Karley

## Abstract

Aphids are significant agricultural pests of cereal crops with a worldwide distribution. The control of aphids in agricultural systems is currently heavily reliant on insecticidal compounds, but it is becoming increasingly apparent that chemical-based control of agricultural pests has far-reaching unintended consequences on agro-ecosystems. As a result, more sustainable means of aphid control are becoming increasingly desirable. Potential options include increasing plant resistance against aphids, promoting biocontrol, and the combined use of both strategies. When used together it is important to understand how, and to what extent, increased plant resistance against aphids affects the success of biocontrol agents. In this current study, we examine how partial-resistance against cereal aphids in a wild relative of barley, *Hordeum spontaneum 5* (Hsp5), affects the success of the common parasitoid of cereal aphids, *Aphidius colemani*. We show that the parasitism success of *A. colemani* attacking nymphs of the bird cherry-oat aphid, *Rhopalosiphum padi*, contained on Hsp5 is reduced compared with the parasitism success of wasps attacking *R. padi* nymphs feeding on a susceptible modern cultivar of barley, *H. vulgare* cv. Concerto. *Explanta* parasitism assays showed that the in parasitoid success is a direct effect of the plant environment (such as differential architectural traits), rather than an indirect effect dur to a decrease in aphid suitability resulting from increased resistance against aphids in Hsp5. Our study highlights the importance of understanding the direct and indirect effects of plant resistance against aphids on biocontrol strategies.

## Introduction

Aphids are major worldwide agricultural pests (VAN EMDEN AND HARRINGTON 2017) and are routinely managed with insecticides. However, due to unintended ecological consequences arising from insecticide use (DEWAR AND FOSTER 2017; GOULSON 2013), more sustainable pest management practices are desirable. Sustainable practices can include the use of aphid natural enemies for biocontrol (RAMSDEN et al. 2017), improving innate plant resistance (ARADOTTIR et al. 2017), and a combination of both strategies (CAI et al. 2009; MITCHELL et al. 2010). When used together, it is important to assess how improved plant resistance might affect the success of natural enemies, especially parasitoid species, as increased plant resistance could reduce aphid viability as parasitoid hosts.

We recently compared a modern domesticated cultivar of barley *Hordeum vulgare* (Linnaeus) cv. Concerto (Concerto) with a wild progenitor species possessing partial-resistance against aphids, *H. spontaneum* 5 (Linnaeus) (Hsp5) (LEYBOURNE et al. 2019). We showed that partial-resistance in Hsp5 primarily involves mesophyll- and phloem-derived resistance factors, associated with increased defence gene expression and a reduction in phloem quality, respectively, which contribute to decreased aphid fitness (LEYBOURNE et al. 2019). However, we have not assessed the extent to which the reduction to aphid fitness, and the altered plant phenotype resulting from the partial-resistance traits, might interact to alter the ability of natural enemies to control aphids.

Here, we examine whether oviposition success of the generalist parasitoid, *Aphidius colemani* Viereck, is differentially affected when attacking the bird cherry-oat aphid, *Rhopalosiphum padi* (Linnaeus), feeding on one of these two plants (susceptible Concerto and partially-resistant Hsp5). We hypothesised that parasitoid oviposition success will be reduced when attacking *R. padi* infesting partially-resistant Hsp5 due to poor aphid performance on this plant. We discuss these results in relation to the unintended consequences of partial-resistance against aphids on higher trophic levels and the potential implications for biocontrol.

## Materials and Methods

### Plant growth and insect rearing

*Hordeum vulgare* (Linnaeus) cv. Concerto (Concerto) and *H. spontaneum* 5 (Linnaeus) (Hsp5) seeds were surface-sterilised by washing in 2% (v/v) hypochlorite and rinsing with d.H2O. Hsp5 seeds were incubated at 4°C for 14 days and Concerto seeds were incubated at room temperature for 48 h, and all seeds were incubated in the dark. Differential seed incubation was required to break seed dormancy in Hsp5 to synchronise germination. Seedlings were planted into compost (Bulrush, Northern Ireland) and grown in randomised blocks until tillering (2.1-2.2 on the ZADOKS et al. (1974) cereal staging key) before use in experiments.

Bird cherry-oat aphids, *Rhopalosiphum padi* (Linnaeus) (genotype B uninfected by defensive endosymbionts; LEYBOURNE et al. (2018)) were reared in ventilated cups on one week old barley seedlings (*H. vulgare* cv. Optic) and *Myzus persicae* (Sulzer) (a mixed population containing genotypes F and O) were reared in ventilated Perspex cages on young oilseed rape, *Brassica napus* (Linnaeus) cv. Mascot. Cultures were maintained at 20 ± 2°C 16:8h (L:D) with PAR 150 µmol/m^−2^ s^−1^. Mummies of the Braconid wasp *Aphidius colemani* (Viereck), supplied by Fargro (West Sussex, UK), were contained in ventilated boxes until eclosion and supplied with 50% (v/v) honey soaked into a cotton wool ball as a food source. A cohort of emerging wasps (5-7 day old) were transferred to *M. persicae*-infested oilseed rape plants enclosed in a fine mesh cage. After 12 days mummies were collected and transferred back to the ventilated plastic boxes until eclosion. To ensure parasitoids had no prior experience of the experimental *R. padi* clones, wasps were reared through at least three generations on *M. persicae* before use in bioassays. Plants and wasp cultures were maintained at 18 ± 2°C and 16h:8 h (day:night).

### Wasp fitness and statistical analysis

Two parasitism assays were performed. For the first assay (*n = 15 for each plant*), plants were infested with 50 1^st^ – 3^rd^ instar *R. padi* nymphs, contained onto plants using microperforated plastic bags. Nymphs were left for 24 h to establish before a single 5-7 day old *A. colemani* female, presumed mated, was introduced. After six hours, the wasp was removed and twelve days later the number of aphid mummies was scored. The second assay (*n = 4 for each plant*) was performed *ex-planta* to disentangle direct and indirect effects of the host plant on aphid susceptibility to mummification by removing the direct influence of the plant structural environment. Briefly, ten *R. padi* nymphs (1st – 3rd instar) were reared on either Hsp5 or Concerto in ventilated cups. Wasp arenas were created in Petri dishes by embedding four barley (cv. Optic) leaves, adaxial surface uppermost, in 1% w/v agar to which the ten *R. padi* nymphs were added. A 5-7 day old *A. colemani* female, presumed mated, was introduced to the arena. After attacki, the nymph was removed and returned to the rearing environment, the wasp was observed until all nymphs had been attacked and returned to the rearing environment. The number of *R. padi* mummies was scored twelve days later and the proportion of the ten nymphs successfully parasitised was calculated. Statistical analyses were carried out using R Studio v.1.0.143 running R v.3.4.3 (R Core Team, 2017) with packages car v.2.1-4 (FOX AND WEISBERG 2011), lme4 v.1.1-13 (BATES et al. 2015), and ggplot2 v.2.2.1 (WICKHAM 2009). For both experimental bioassays, linear mixed regression models were used to model *A. colemani* fitness (the proportion of *R. padi* mummified) in response to plant type, with experimental block as a random model factor. X^2^ Analysis of Deviance test was carried out on the final models.

## Results and Discussion

In line with our predicted hypothesis, the oviposition success (proportion of *R. padi* successfully parasitised) of *A. colemani* was reduced when attacking *R. padi* infesting the partially-resistant wild relative of barley, Hsp5, compared with *A. colemani* attacking *R. padi* infesting the susceptible cultivar Concerto (X^2^_1_ = 7.59; p = 0.005; Fig. 1). Previous studies have reported similar decreases in parasitoid wasp fitness when attacking aphids infesting plants with increased resistance (CHACÓN et al. 2012; KALULE AND WRIGHT 2007; MITCHELL et al. 2010; ODE 2006; ODE AND CROMPTON 2013; SCHÄDLER et al. 2010), although this is not often associated with specific plant traits (ODE 2006). As we have previously characterised the mechanism of partial-resistance in Hsp5 (LEYBOURNE et al. 2019) we can predict how these traits might have affected parasitoid success. The partial-resistance mechanism of Hsp5 involves elevated defence gene expression and reduced phloem nutritional quality, resulting in decreased aphid fecundity and nymph mass in three cereal aphid species (LEYBOURNE et al. 2019) and several *R. padi* genotypes (LEYBOURNE et al. 2018). The primary effect of these factors would be to decrease aphid quality as viable hosts for *A. colemani* due to reduced nymph quality and size.

**Fig. 1:**
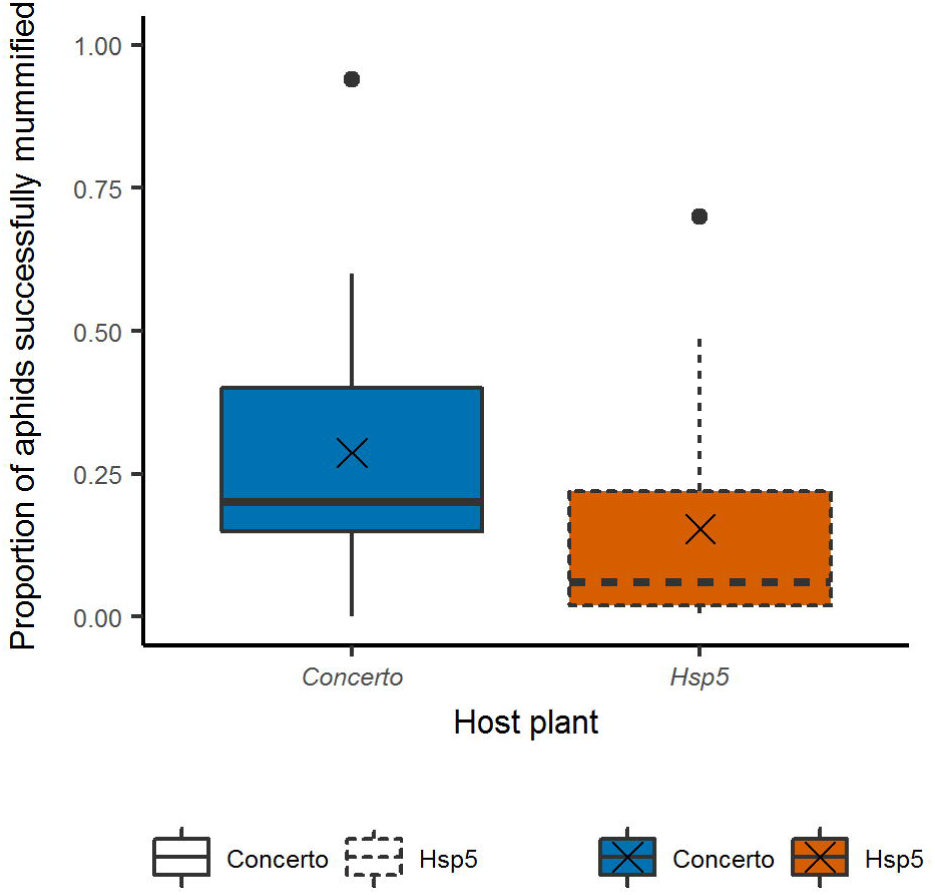
The proportion of *Rhopalosiphum padi* nymphs successfully mummified by *Aphidius colemani* (median and inter quartile ranges) when nymphs were infesting either the susceptible barley cultivar (*H. vulgare* cv. Concerto) or the partially-resistant wild relative (*H. spontaneum* 5 - Hsp5). *n = 15* for each plant. The cross (“x”) marks the mean value.

To examine whether reduced aphid quality was the primary cause of the decrease in *A. colemani* oviposition success on Hsp5 compared with Concerto, rather than a direct effect of differential plant traits, *ex-planta* assays were carried out. These assays showed that the proportion of mummified *R. padi* did not vary with aphid rearing environment (Concerto vs. Hsp5: X^2^_1_ = 0.06; p = 0.796; Fig. 2). This indicates that the observed decrease in *A. colemani* oviposition success on Hsp5 (Fig. 1) is likely a direct result of the plant environment rather than a decrease in aphid suitability as a host. Hsp5 has a higher density of non-glandular leaf trichomes and a less diverse epicuticular surface chemistry compared with Concerto (LEYBOURNE et al. 2019). An increased abundance of non-glandular trichomes could lead to a reduction in oviposition success by acting as a physical barrier and inhibiting the searching efficiency of *A. colemani*. Indeed, computational models have shown that parasitoid success decreases on plants with a more complex physical structure (GINGRAS et al. 2002). It is important to note that other plant traits which have not yet been characterised might have also influenced parasitoid searching and oviposition success. These could include architectural traits, such as plant compaction (PRADO AND FRANK 2013) and leaf wax bloom (CHANG et al. 2004), and differential emission of plant volatiles between the two plant types, which could be attractive (MC CORMICK et al. 2012) or repellent (SNOEREN et al. 2010) to parasitoids. It is also possible that aphids feeding on Hsp5 were less settled than aphids feeding on Concerto, leading to an increase in dropping in response to A. colemani presence, which could influence parasitoid success (CHAU AND MACKAUER 1997).

**Fig. 2:**
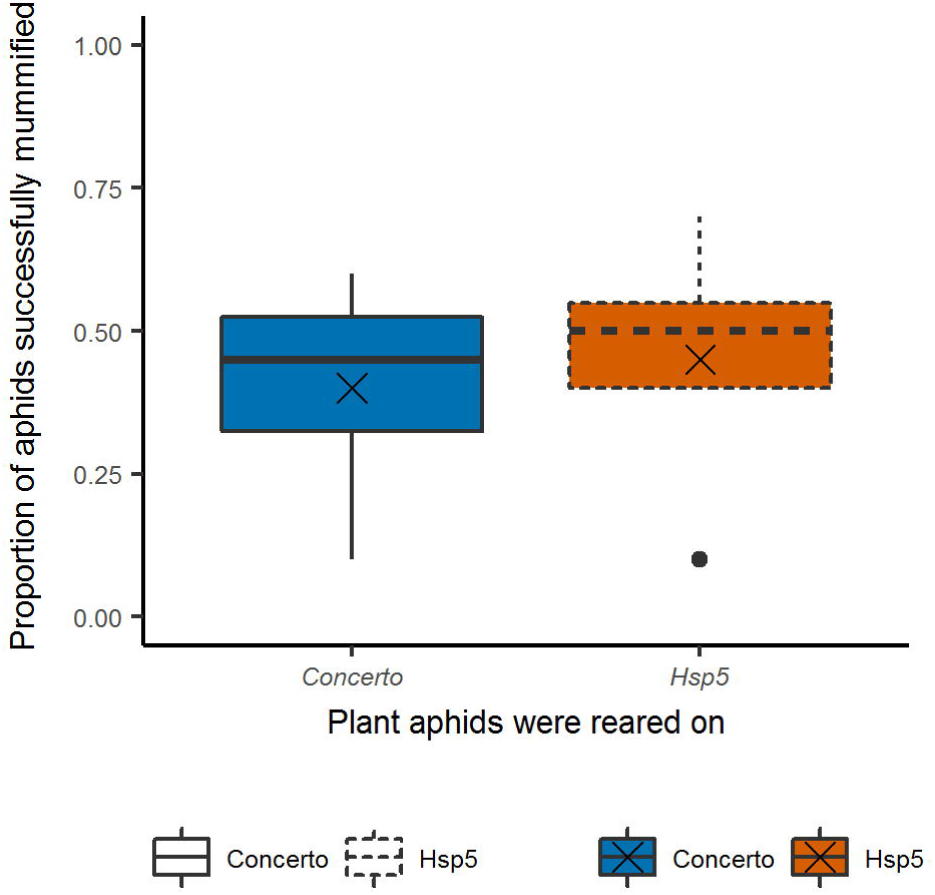
The proportion of *Rhopalosiphum padi* nymphs successfully mummified by *Aphidius colemani* (median and inter quartile ranges) in *ex-planta* assays when aphids had been reared on either the susceptible barley cultivar (*H. vulgare* cv. Concerto) or the partially-resistant wild relative (*H. spontaneum* 5 - Hsp5). *n = 4* for each cultivar. The cross (“x”) marks the mean value.

Our results show that plant partial-resistance against cereal aphids can have negative impacts on natural enemy success, in line with findings from other plant-aphid-parasitoid systems (KALULE AND WRIGHT 2007; MITCHELL et al. 2010; ODE 2006). We show that this reduction in parasitoid success is not a direct consequence of decreased nymph suitability as a host, as we hypothesised, but instead is potentially associated with indirect effects of the plant environment on parasitoid oviposition behaviour. We suggest that this could be driven by differences in plant architecture between the two plant types, including non-glandular leaf trichome density. Our study highlights the importance of understanding the direct and indirect effects of plant resistance against aphids on biocontrol agents.

## Acknowledgements

D.J.L was funded by the James Hutton Institute and the Universities of Aberdeen and Dundee through a Scottish Food Security Alliance (Crops) PhD studentship. A.J.K. and T.A.V. were funded through the strategic research program funded by the Scottish Government’s Rural and Environment Science and Analytical Services Division. J.I.B.B was supported by the European Research Council (310190-APHIDHOST). We would like to extent thanks to Carolyn Mitchell (James Hutton Institute) for helpful comments on the manuscript.

## Author contributions

D.J.L and A.J.K conceived the study. D.J.L carried out the experimental work. D.J.L analysed the data and conducted statistical analyses. All authors contributed to data interpretation. D.J.L wrote the manuscript with input from all authors. J.I.B.B. and A.J.K secured funding. All authors read and approved the final manuscript.

## Data availability

Data and statistical code are available from the corresponding author upon reasonable request.

